# Integrated multi-platform approaches to gain insights into ecosystem’s fundamental ecology and habitat specific alterations

**DOI:** 10.1101/2024.06.07.597995

**Authors:** Shekhar Nagar, Chandni Talwar, Ram Krishan Negi

## Abstract

The increasing availability of metagenome-assembled genomes and environmental metagenomes provides unprecedented access to the metabolic potential and functional differences within the habitats. The hot spring microbiome with its diverse habitats and relatively well-characterized microbial inhabitants offers an opportunity to investigate core and habitat specific community structures at an ecosystem scale. Here, we employed tailored genome-resolved metagenomics and a novel approach that offers metagenomic overlaps to investigate the core and habitat-specific microbial diversity and multifunctionality of microbial residents of three habitats: microbial mat, sediment and water. We recovered 6% of the Ecosystem core community (ECC) in the habitats suggesting the widespread acquisition of Proteobacteria involving in the diverging trajectories of the hot spring and 72% of the Habitat specific community (HSC) in microbial mat, sediment and water habitats offers insights into specific adaptations due to extreme conditions. Strain-level resolution of metagenome-assembled genomes defined the habitat specific genotypes (HSGs) and comparative metagenomic analysis exposed ecosystem-core genotypes (ECGs). Further, the functional attributes of ECGs revealed a complete metabolic potential of nitrate reduction, ammonia assimilation and sulfate reduction. The highest cycling entropy scores (H’) of N cycle suggested the enrichment of nitrogen fixing microbes commonly present in all three habitats. While specifically HSGs possessed the amino acid transport and metabolism functions in microbial mat (9.5%) and water (13%) and 19% of translation, ribosomal structure and biogenesis in sediment. Our findings provide insights into population structure and multifunctionality in the different habitats of hot spring and form specific hypotheses about habitat adaptation. The results illustrated the supremacy of using genome-resolved metagenomics and ecosystem core metagenomics postulating the differential ecological functions rather than that of explaining the presence of functions within ecosystem.

## Introduction

For centuries ecologists have studies how the microbial diversity interacting and existing in various biomes. They have begun the exploration of single habitat microbial communities considering soil habitat as central hotspot for biogeochemical cycles and major consortium of living biomass in terrestrial ecosystems (Whitman et al., 1998, Lehmann et al., 2017, Philippot et al., 2024). Different habitat typically harbor distinct assemblages of microbial taxa and and their geochemical functions in corresponding confined biogeographical area. The increasing availability of microbial repertoire and environmental shotgun metagenomes provide insights into the overlapping microbial hubs present in habitats with a tremendous significance to fix the atmospheric uncertainties.

Ecosystem is a fundamental organizational unit of the biosphere in which biological communities interact with their non-biological environment through energy flows and material cycles (Odum, 1971). Ecosystem functional ecology could be studied using the micro-organisms which derives the patterns, processes and services of ecosystem. The functional ecology attributes to the ecological function concept the aim of explaining ecosystem processes rather than that of explaining the presence of organisms within ecosystem (Dussault, 2019). The stochastic events (random changes) of ecosystem are majorly account for population interaction and habitat selection. Recent studies have focused on underlying mechanisms that shape the biogeographic patterns among most free-living microbial taxa (Wang *et al.,* 2017). There are some evidences that these microbes are also capable of altering and conducive to the cycling of essential nutrients (Schimel and Schaeffer 2012; Seto and Iwasha, 2020) The parameters such as geographical distance, pH, temperature, vegetation and aridity etc roles supports the growing evidence of structuring microbial community (Chu *et al.,* 2010; Maestre *et al.,* 2015; Prober *et al.,* 2015). The stochastic events are the reason behind species abundance and diversity while neutral theory suggests that all individuals in the community are ecologically equivalent (Hubbell, 2001). These events led the dispersal of similar individuals to nearby sites or closer sites with similar environments. The dispersal of microbes could be specific or random depending upon the ability of the hosting microbes studied in different habitats such as desert (Ronca *et al.,* 2015), permafrost (Yergeau *et al.,* 2007), aquatic ecosystems (Zinger *et al.,* 2014). Environmental microbiome research has demonstrated numerous factors including biogeochemical cycles, potential pathogens, essential microorganisms and human health (Bolan and Hedley, 2003; Husson O 2013; Koch and Dahl, 2018, Sokol et al., 2022).The branch of bioinformatics have become more focused on sequencing errors and high coverage assemblies. Yet, no studies up to date have focused on providing insights in the communal and specific genetic repertoire across different habitats of a hot spring.

The hot spring microbiome is an ideal system in which to investigate microbial population structures in a complex lamdscape. Different microbes in terrestrial niches such as water, microbial mat, sediments needs to be explored to understand the third dimensional concepts of ecosystem. Although our understanding of the genome networking of these niches continue to expand, and there has been inadequate progress in understanding how the functional proficiencies of these niche specialized communities varies across biomes. To investigate the core community dynamics and the mechanisms underlying bacterial communities among habitat types (microbial mat, sediment and water), we conducted genome-resolved metagenomics and a comparative metagenomic analysis of a mesothermic hot spring Khirganga atop Northern Himalayas (Nagar *et al.,* 2022). These approaches uncovered a metabolic overlap and differentiation of three habitat types where the abiotic factors like pH, temperature and altitude remain constant. We attempted to address the following (i) 16SrRNA community differentiation (ii) strain level resolution of the reconstructed genomes from the hot spring which confers as habitat-specific attributes (iii) computed orthologous segments of the habitat types termed as ecosystem core genetic repertoire. The study described a novel comparative metagenomic approach i.e., ecosystem core metagenomics and *denovo* extraction of strains from metagenomes to evaluate ecosystem functions.

## Materials and methods

### Statistical Analysis

The statistical analyses were performed using the SPSS 17.0 software package (SPSS Inc., Chicago, USA). Normality of all weight and length data of fish were checked by Shapiro-Wilk’s test. Values of the taxonomic compositions (except grouped abundances) were expressed as the mean ± SE. Significance of the differential abundances of taxa was determined using a two-tailed t-test on grouped taxonomic abundances (significance declared at *P* <0.05). Statistical analysis of shared taxonomic diversity across habitats was performed using the Statistical Analysis of Metagenomics Profiles (STAMP) program (http://kiwi.cs.dal.ca/Software/STAMP), retaining unclassified reads (Parks *et al.,* 2010). P-values were calculated by Welch’s two-sided t-test and subjected to multiple testing correction using the Benjamini-Hochberg method for a False Discovery Rate (FDR) of 1% (Benjamini and Hochberg, 1995). The consensus profile level were automated in order to correlate the different habitat profiles in a pairwise fashion followed by the parent level.

### Identification of Habitat specific communities (HSC) and Ecosystem core communities (ECC)

Raw reads were subjected to trimming and quality filtering by DADA2 (Callahan *et al.,* 2016). The representative sequences thus obtained were then again filtered in the next step to only contain sequences displaying hits against the reference Greengenes database [Greengenes OTUs (16S) 13_8] with >99% confidence level (DeSantis *et al.,* 2006). The extracted reads were assembled *de novo* with REAGO (Yuan *et al.,* 2015) followed by clustering at 97% similarity using UCLUST integrated in QIIME (Caporaso *et al.,* 2010). The clustered ‘Operational taxonomic Units’ (OTUs) were then assigned to representative genera and species using RDP naïve Bayesian classifier (Cole *et al.,* 2007). OTUs were clustered if they shared at least 97% similarity with the database sequences at a percentage query length alignment threshold of 95%. In the final step, OTUs derived from eukaryota, chloroplast and mitochondria were removed to generate an OTU table and diversity was analyzed using QIIME 2 (Bolyen *et al.,* 2019). Sampling and sequencing depth were studied through rarefaction curves constructed by plotting species richness as a function of number of sequences. Percent composition was calculated as the percentage of taxa in a given phylum relative to the total number of taxa. The OTUs were aligned using CLUSTALW and phylogeny was constructed using Maximum-Parsimony algorithm using MEGA v7.0 (Kumar *et al.,* 2016). Bootstrap tests were performed using 500 resamplings. Diversity at different hierarchical levels was plotted for comparison between two groups of samples based on OTU abundances using the “metacoder” package in “R” (Foster *et al.,* 2017, R Development Core Team, 2015). To determine taxa with significantly enriched abundance in either of the three community types, we applied a two-tailed t-test on grouped taxonomic abundances implemented within QIIME (P⩽0.05 deemed significant) and analogous comparisons were made.

We utilized a standard approach with G-test of Independence analysis to reconstruct the habitat-habitat networks with 16S rRNA datasets of microbial mat, sediment and water (Ma *et al.,* 2015, 2016). A G-test for independence (McDonald, 2015) was used to test whether sample-nodes within categories are more connected within a group than expected by chance. Further, the OTUs were partitioned into Habitat specific communities (HSC) and Ecosystem core communites (ECC) to create a network of distinct as well as OTUs shared among habitats using QIIME (Caporaso *et al.,* 2010). The network was visualized in Cytoscape v3.7.1 (Shannon *et al.,* 2003) and OTU connections were statistically analyzed to provide support for the clustering patterns.. Further, phylogeny of the OTUs was recapitulated by constructing a dendrogram in iTOL v3 (Letunic and Bork, 2016).

### *In-silico* strategies for identification of Ecosystem homologues and Habitat specific genetic repertoire

In order to access the common genetic repertoire across the habitats through ecosystem core-metagenomics, a cross-assembly was achieved using MEGAHIT (Li *et al.,* 2015) and contigs shorter than 1,000 nts were discarded from further processing. To estimate the coverage of each contig in each sample, a contig coverage database was generated reads were mapped on to the assembly using Bowtie2 (Langmead and Salzberg, 2012) and sorting and indexing were performed over the binary outputs using samtools (Li *et al.,* 2009). Further the contigs were scanned for commonly set of archaeal and bacterial single copy genes using Hmmer v3.3.1 (Eddy SR, 1998) to calculate redundancy and completeness. Contigs were clustered according to the tetra nucleotide frequency and GC content using CONCOCT v2 (Alneberg *et al.,* 2014) and taxonomy from Centrifuge Archaeal Bacterial database 2018 (Kim *et al.,* 2016) was defined in anvi’o (Eren *et al.,* 2015).

In order to gain insights into habitat-specific functional potential, we tried to investigate the metagenome-assembled genomes (MAGs) reconstructed from three different habitats of the hot spring (Nagar et al., 2023). The homologues gene content of all 39 bacterial MAGs were estimated using microbial pangenomics workflow in anvi’o (Eren *et al.,* 2015) and based on the distribution of gene clusters, the genomes were partitioned using MCL algorithm into core and accessory (habitat specific) gene content (Distance: Euclidean; Linkage: Ward). The accessory gene sequences of each individual MAG was further introduced in Denovo Extraction of Strains from Metagenomes (DESMAN) to infer the variable positions of accessoy genes (Quince *et al.,* 2017). The accessory gene sequences of each individual MAG were optimized in DESMAN to infer the variable positions of the species core gene content. Here, the strain variation was resolved by identifying the core genes with the reference genomes or related taxa that were available in database, these identified orthologs were designated as single-copy core species genes (SCSGs). By inferring the COGs of SCSGs, we were able to estimate variant positions of alleles for every single genome using likelihood ratio test applied to the frequencies of each base summed across genomes. These unique sequences and haplotypes of each core gene was unified. Altogether, the resulted strain-specific gene content of individual MAGs with similar habitat were incorporated in anvi’o so as to determine the core gene clusters of the individual habitats denoted as habitat specific genotypes (HSGs). Summarizing these strain resolved genotypes habitatwise allowed us to detect and investigate the frequencies of habitat specific genes (HS_g) among populations. The HSGs were also obtained from ecosystem core-metagenomic binning methods where non-homologues segments and their encoded genes were termed as HSG_g. The schematic representation of the integrated approaches were shown in Figure 1.

**Figure 1:**
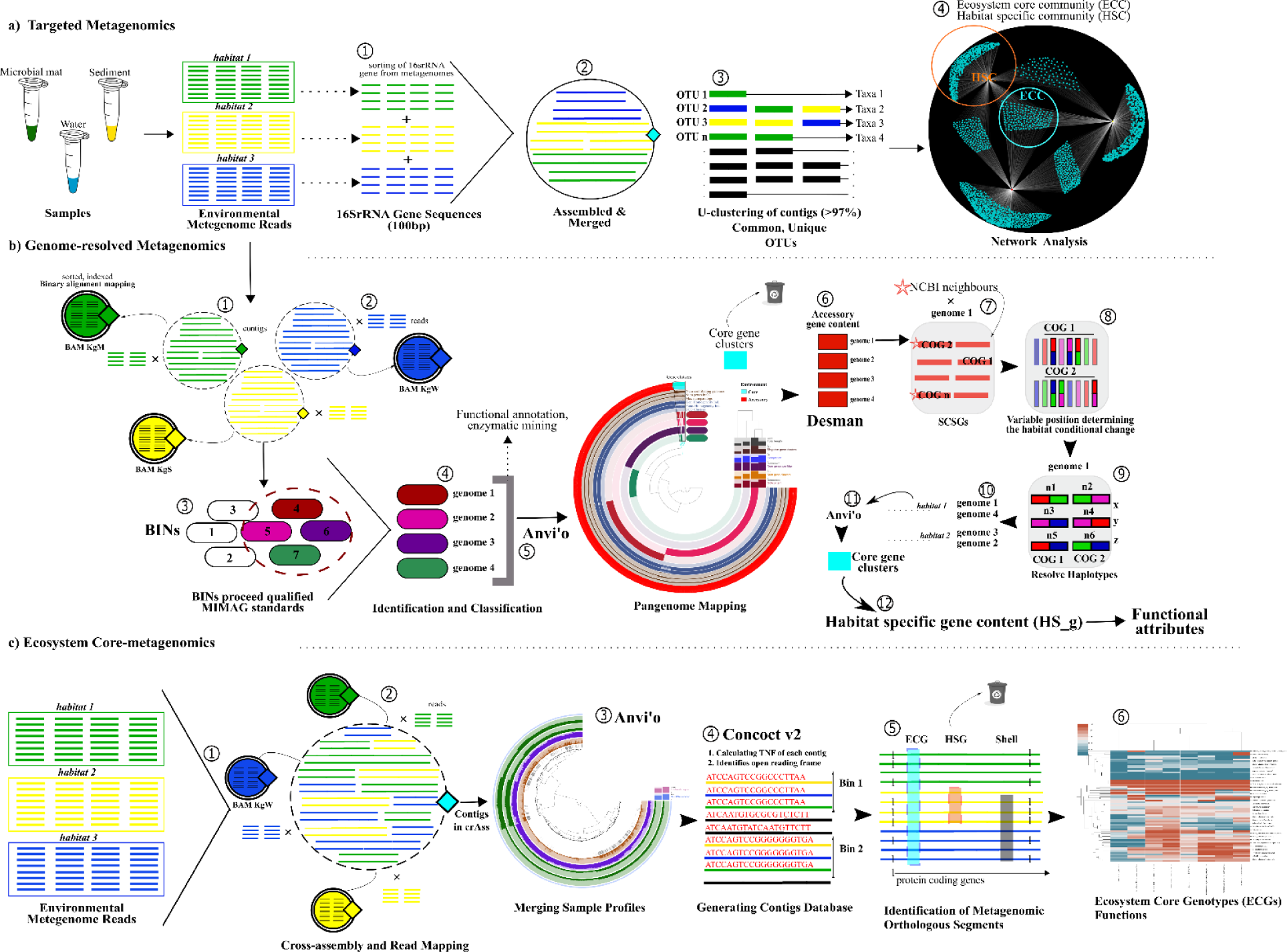
Schematic representation to estimate the ecosystem core and habitat specific diversity and functions. **a) Targeted metagenomics:** (1) sorting of the 16S rRNA reads from paired-end sequences of microbial mat (green), sediment (yellow) and water (blue) environmental metagenomes, (2) assembled and merged together, (3) clustering of contigs (cut-off >97%) to examine Operational Taxonomic Units (OTUs) and then assigned to representative taxa, (4) network analysis using G-test of independence (p-value < 0.001) and clustered groups were labelled as Habitat-specific community (HSC) and Ecosystem core community (ECC); **b) Genome-resolved metagenomics to determine Habitat specific gene content (HS_g):** (1), (2), (3), (4) discussed in objective III methodology section, (5) pangenome mapping of genomes (39 bacterial, 12 phyla) was done and (6) claimed accessory gene content of each genome was separately analysed using DESMAN (7) identification of Single core-species genes (SCSGs) in MAGs (8) calculating variable positions by mapping reads to contigs associated with SCSGs of each genome, (9) haplotype deconvolution was achieved using 500 iterations (10) Each genome was aligned with their reconstructed habitat-based individdual in order to determine HS_g (11) HS_g bins were exported from anvi’o orthologous rule (12) Functional annotation of the HS_gM (microbial mat), HS_gS (sediment) and HS_gM (water) were estimated; **(c) Ecosystem core-metagenomics** (1) Cross-assembly of the reads for all the six samples (2) coverage, mapping of reads over the contigs and profiling was completed (3) profiles of all six samples were merged in anvi’o (4) binning process was accomplished using Concoct v2 (5) common bins (orthologous segments) were obtained and taxonomically characterized (6) common genetic repertoire was integrated in MEBS to resolve significant biogeochemical cycles of ecosystem

The gene clusters of ecosystem core (EC_g) and habitat specific (HSG_g, HS_g) principles were further classified into ORFs using Prodigal v2.6.3 (Hyatt *et al.,* 2010). Further, the HS_g was annotated from NCBI COGs database (Galperin *et al.,* 2019) using rpsblast v2.2.5, and also searched using hmmscan v3.0 (Eddy SR, 1998) with a prediction cut off e-value 0.001 over Pfam v24.0 database (Finn *et al.,* 2010) and reprsented on the basis of the abundance of proteins with contribution of each genome. Whereas, the normalized Pfam entropy scores of essential elements Nitrogen (N), Sulfur (S), Iron (Fe), Oxygen (O) and Carbon (C), and their clustered metabolic proteins completeness in EC_g and HSG_g were computed using Multigenomic Entropy Based Score (De Anda *et al.,* 2017).

## Results and Discussion

### Statistical Analysis

No significant differences in diversity profiles were observed within the two metagenomes of each habitat (paired sample t-test, p-value ≥0.05). There is no significant difference between the two replicates of each sample i.e. p-value is greater than 0.05 in all three combinations. The metagenomic profiling among three habitats attained similar taxonomy profiling with respect to inter-class level taxonomy (Figure S1). Statistically, after filtering with probability (p ≥0.05) the class-taxonomy profiling of the datasets shows correlation among all three datasets with correlated values i.e. microbial mat and sediment (r^2^ = 0.542), water and sediment (r^2^ = 0.912) and microbial mat and water (r^2^= 0.711). The water and sediment showed a positive linear correlation coefficient with class abundance profiles may be shaped by equivalent environmental parameters (Zhang et al., 2021). While water habitat abundance profiles also showed a moderate positive linear correlation with microbial mat suggested that the microbial communities were not differently structured due to identical physiochemical parameters irrespective of different habitats (Fray et al., 2024).

### 16SrRNA abundant profiles

Diversity analysis based on reconstructed 16S rRNA sequences from short length reads and their taxonomic clustering resulted in a total of 2688 OTUs with ≥ 97% sequence similarity of 35 bacterial phyla and three archaeal phyla across all the three datasets. Of these, 81.54% of the OTUs could be assigned to bacterial or archaeal (0.5%) sequences (Figure 2A). The remaining 18.46% of the reads that remained unclassified reflect at the incompleteness of the 16S rRNA databases and a high untapped microbial diversity at extreme habitats as are the hotsprings. Among three datasets, water had greater richness and diversity but were not significantly different than microbial mat and sediment sets. *Proteobacteria* were found to be the most dominant phylum (44.60%) followed by *Chloroflexi* (10%), *Bacteroidetes* (9.75%), *Planctomycetes* (5.52%), *Cyanobacteria* (5.15%), *Firmicutes* (5.09%), *Spirochaetes* (3.14%) and *Acidobacteria* (2.14%) comprising about 90% of the 16S rRNA in all three of the samples (Figure 2B). Members of these phyla are commonly observed in thermal springs around the world (Hall *et al.,* 2008; Pagaling *et al.,* 2012; Bowen de León *et al.,* 2013; Wang *et al.,* 2013). The copy number of 16S rRNA gene was mapped along the different taxa which showed that the maximum no. of these genes in microbial mat (n= 180, 77), sediment (n=580, 198), water (n=740, 300), belongs to phylum *Proteobacteria* and class *Gammaproteobacteria* in microbial mat and *Alphaproteobacteria* in sediment and water. Moreover, Archaea was absent in water (Figure 2B). The phylogenetic relationships between the 16s defined OTUs showed an uneven placements on the cladogram suggested the typical assemblage of the microbiota in three habitats (Figure S2).

**Figure 2:**
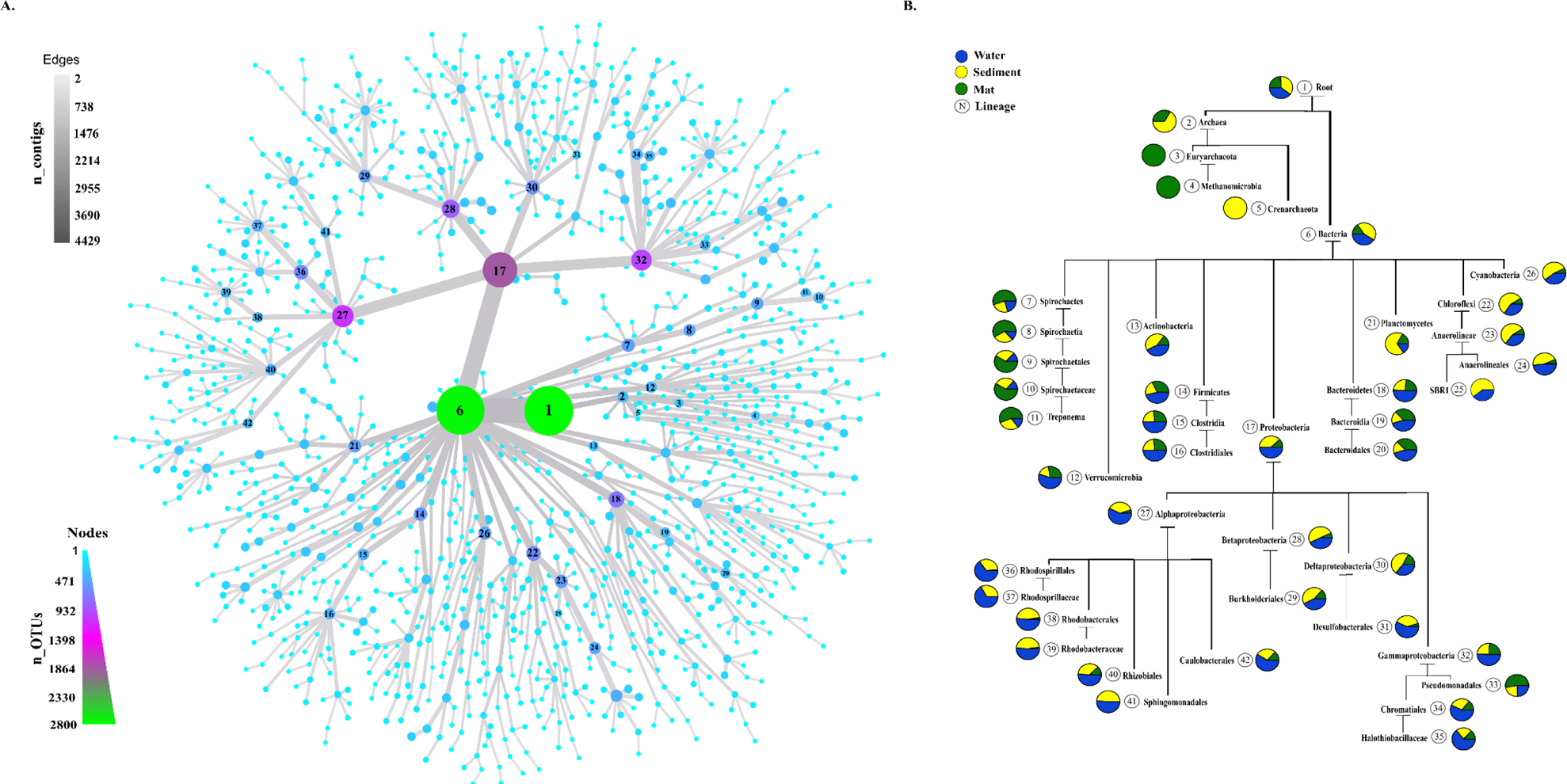
Bacterial and archaeal abundance at different taxonomic hierarchial levels based on 16S rRNA gene sequences. A taxonomic tree constructed from clustered OTUs identified through RDP Naïve Bayesian classifier v2.11. Size and color of nodes and edges are correlated with the abundance of organisms in each community. Out of total 18 bacterial phyla, *Proteobacteria, Bacteroidetes, Spirochaetes, Acidobacteria, Chloroflexi, Firmicutes and Planctomycetes* were dominant in the overall microbial diversity (relative abundance >0.8%). Among archaeal phlya: *Euryarchaeota* and *Crenarchaeota* (class: *Thaumarchaeota*) were the most abundant in all habitat.

### Network analaysis reveal Ecosystem Core and Habitat Specific Community designation

An OTUs network analysis employed to reveal habitat specific community (HSC) and ecosystem core community (ECC) using 16S rRNA and bins generated by clustering contigs on the basis of a combination of hierarchical clustering and taxonomic identity. The network represented a total of 2691 nodes (MSW, n=3) connected to a single component and with 3595 edges. Accordingly, the habitat-habitat interactions provide the percentage ratio of HSC out of total 2688 OTUs i.e., 514 (19.12%), 709 (26.37%) and 748 (27.48%) observed in microbial mat (M), sediment (S) and water (W) samples respectively (Figure 3A). Excluding the unassigned OTUs, the maximum percentage HSC in all three samples were dominated by *Proteobacteria* (27.4 ± 9.6), *Bacteroidetes* (7.2 ± 4.0)*, Chloroflexi* (5.7 ± 3.2)*, Planctomycetes* (3.5 ± 2.6)*, Chlorobi* (3.3 ± 1.3), *Firmicutes* (3.0 ± 0.8)*, Spirochaetes* (2.0 ± 1.3) and *Cyanobacteria* (1.4 ± 0.8).

**Figure 3:**
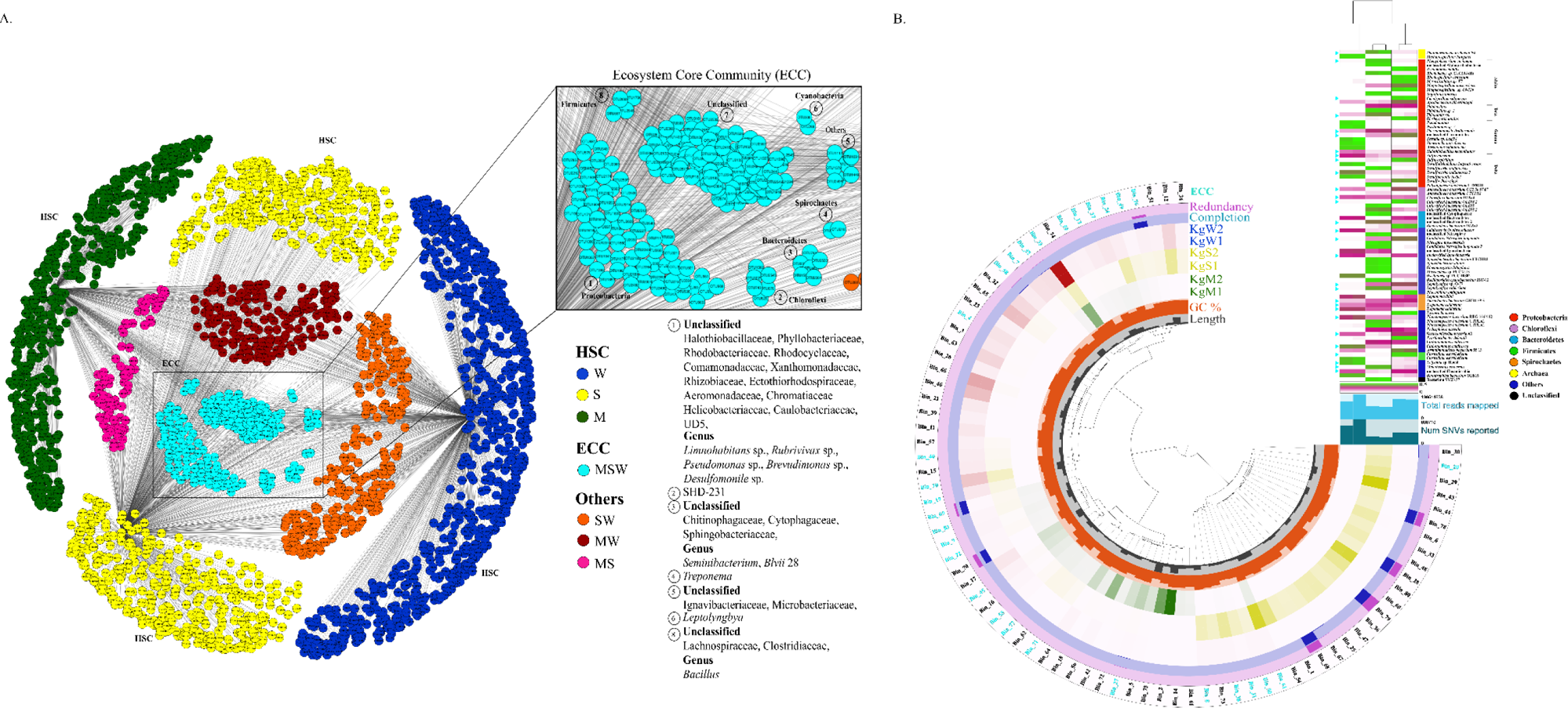
16S rRNA genes abundance inter-habitat interactions and whole metagenome overlaps between microbial mat, sediment and water. (A) Network representation formed through mapping M (microbial mat), S (sediment) and W (water) nodes (OTUs) together to derive out the inter-habitat OTU relationship. The network clearly represents the diversity and clustering on the basis of the relatedness within common (ECCs) and distinct nodes (HSCs). (B) The metagenome contigs are mapped to depict the overlaps of common (ECGs) and habitat specific bins (HSGs) and taxa, abundance of each bin is shown with the heatmap and strip color, respectively.

While, ECC in all three samples (MSW) were 6.51% (175) of the total OTUs identified, percentage dominated by the P*roteobacteria* (17.7%), followed by *Chloroflexi* (8%)*, Bacteroidetes* (6.2%)*, Cyanobacteria* (4%)*, Firmicutes* (3.4%) and *Spirochaetes* (3.4%) (Fig 4A). Also, the water and sediment samples (SW) shared 248 OTUs (9.23%), while water and microbial mat samples shared 197 OTUs (7.32%) and 106 (3.9%) of all OTUs were shared within microbial mat and sediment samples (Figure 3A). The abundance of HSC across all the habitats suggested that hot spring have strong nich specialization irrespective of the overlapping ranges of abiotic factors.

**Figure 4:**
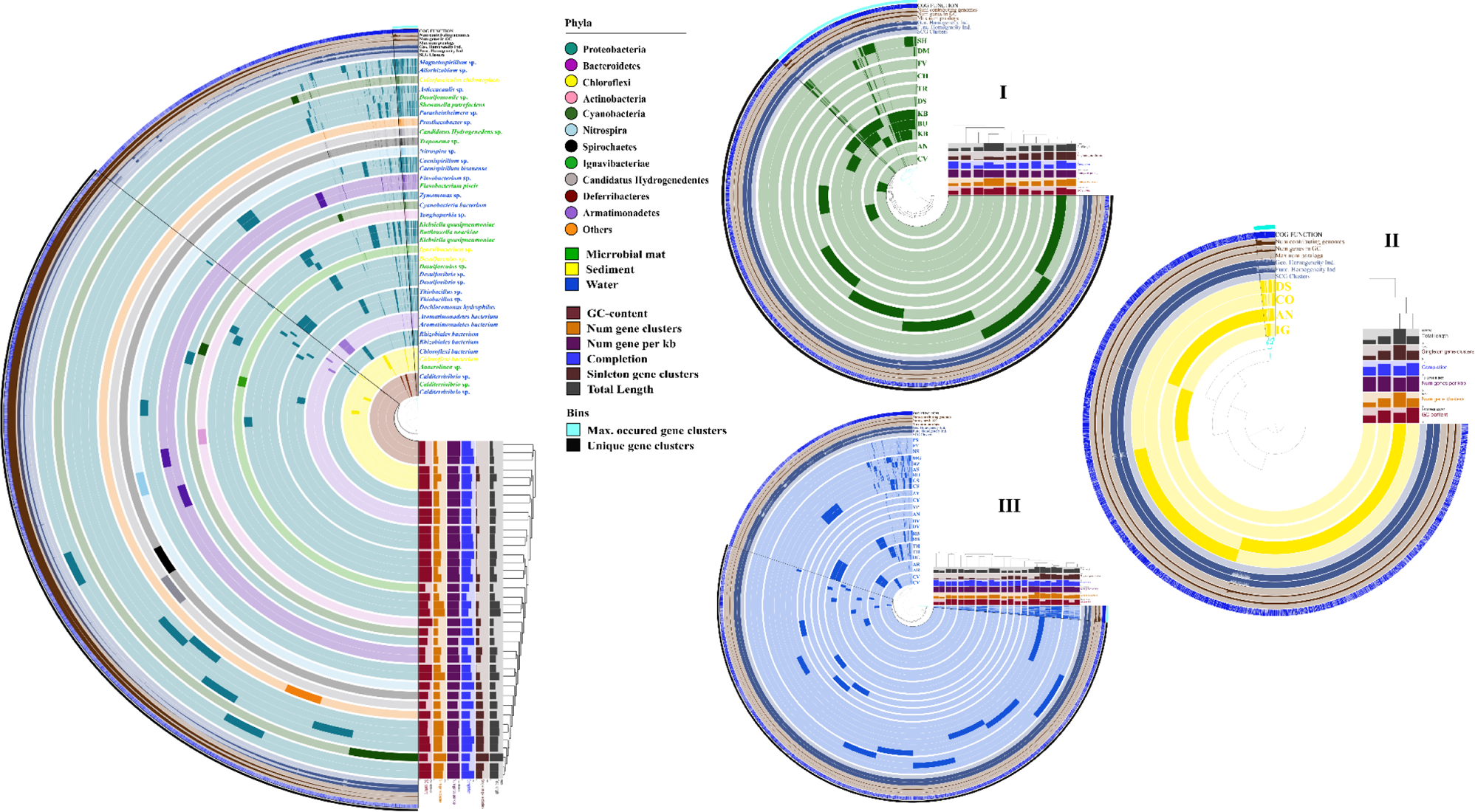
Determination of habitat-specific gene clusters (HS_g). The inner radial shows the 105,589 gene clusters in the pangenome, clustered by presence/absence [Euclidean distance; Ward linkage] across genomes. The 39 bacterial genomes are plotted on the innermost 39 layers (180^ᵒ^ arcs), spaced to reflect discernable clades based on genome abundances. The colors of phyla showed in the middle legends. The outermost seven layers with 2 bins (max. occurred and unique) are colored according to the presence of gene (atleast in one genome), (**I)** bacterial genomes in 11 layers (270^ᵒ^ arcs) reconstructed from microbial mat after strain resolution mapped, showed here maximum occurred common gene clusters (∼11,000) **(II)** 4 genomes in sediment samples shows 773 gene clusters **(III)** 12 genomes in water samples were showing a core gene cluster value of 7923 with a maximum differences among 6 phyla.

### Ecosytem core and Habitat specific gene content analysis

A comparative metagenome approach provides an integrated overview of how the genes are distributed commonly across ecosystem. A conceptual schematic respresntation of ecosystem core-metagenomics, described how we used and integrated first time this naïve approach to combine three different metagenomes samples of hot spring in Figure 1. Cross-assembling the metagenomes (contigs, n= 648,416) provide pertinent observations regarding the regulatory functional genes enrichment commonly in microbial mat, sediment and water. The ecosystem core-metagenomics was used to reveal the orthologous segments (bins) across the three habitats which were denoted as ecosystem core genotypes (ECGs). The presence and absence of gene clusters in individual metagenome could be displayed in anvi’o so that sets of homologous genes are easily identified that are shared by all metagenomes (Figure 3B). Initially, the merged profile database that was generated contained 192,427 ECG contigs which correspond to 29% of all contigs and 69% of all nucleotides (2.23Gb) found in contig database. Total 29 ECGs were determined from the cross-assembled contigs belonged mainly to *Proteobacteria* (n = 9; Figure 3B). The other ECGs were classified into 14 different archaeal and bacterial phyla viz., *Chloroflexi, Spirochaetes* (n= 3), *Cyanobacteria* (n= 2), *Thaumarchaeota, Deferribacteres, Nitrospirae, Chlorobi, Planctomycetes, Verrucomicrobia, Armatimonadetes, Firmicutes, Actinobacteria, Deinococcus-Thermus, Elusimicrobia* (n= 1). The major abundance of taxa in every single ECGs were enlisted in Table 1. The species *Chloroflexi bacterium* OLB14, *Treponema caldarium* (n= 2; Table) together with 25 ECGs were continually communicated in all three habitats of hot spring. A large set of EC_g (genes) recovered from these ECGs were comprising of 38.8% (n= 554,800) of the total pangenome (N= 1,459,679) correspond to complex habitat metagenomes. These results point towards the existence of apparent generalist populations structured together in distinct habitats of hot springs. ECGs could reveal the inter-habitat gene transitions in the populations over a course of time. The genotypes that were recovered were not complete genomes instead they were announced to represent diversity and a composite functional assessment of the ecosystem.

**Table 1:**
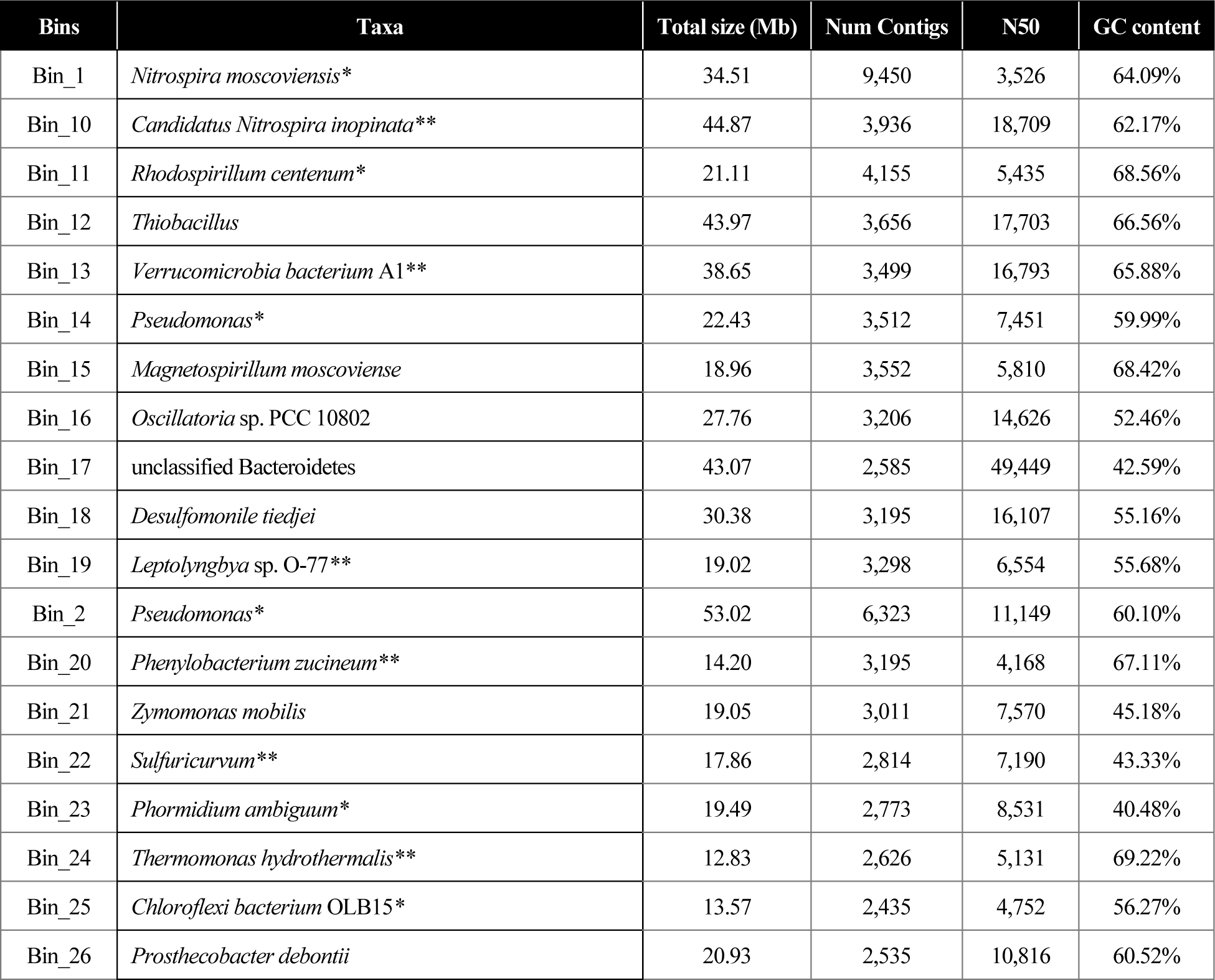
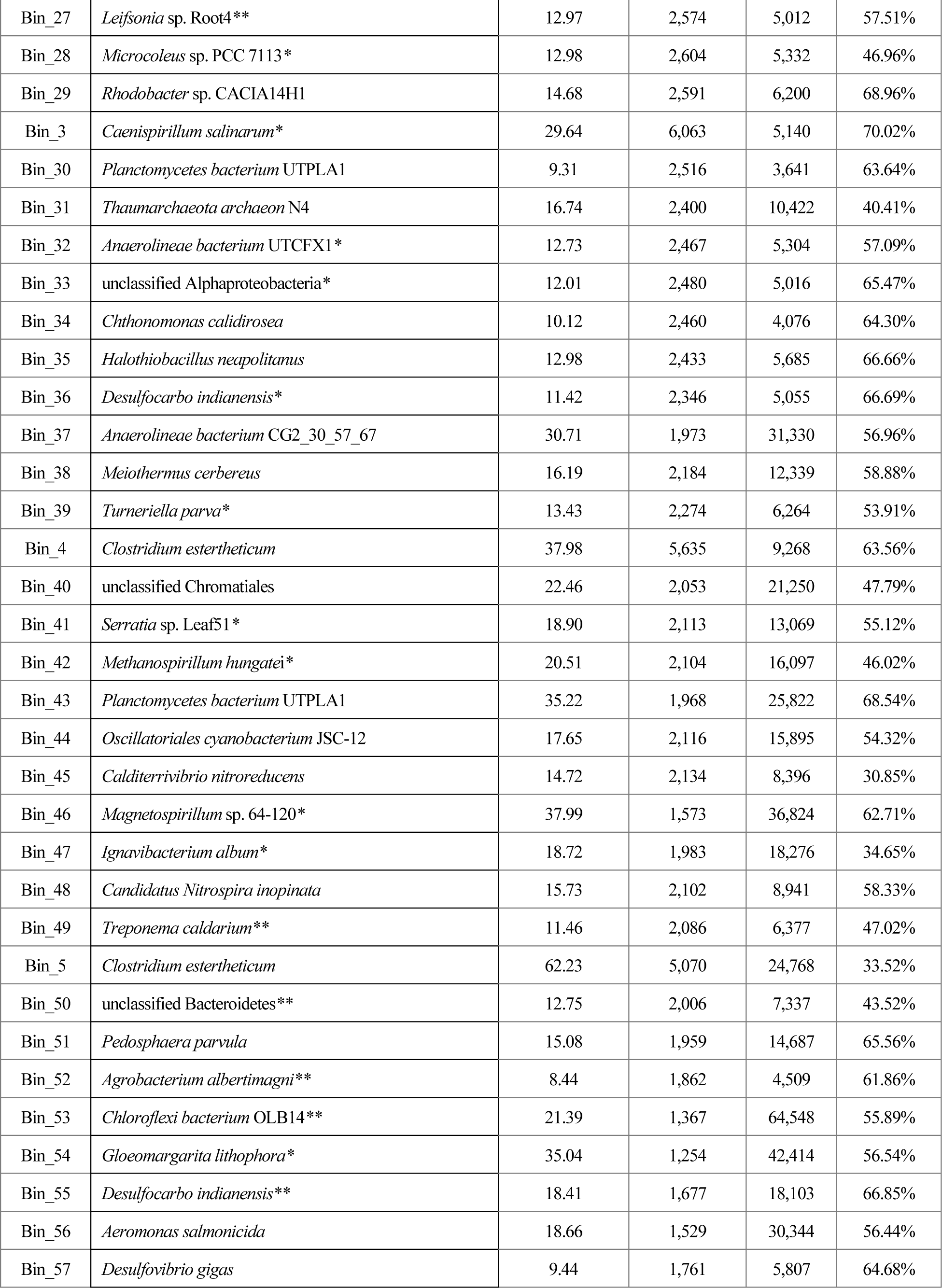
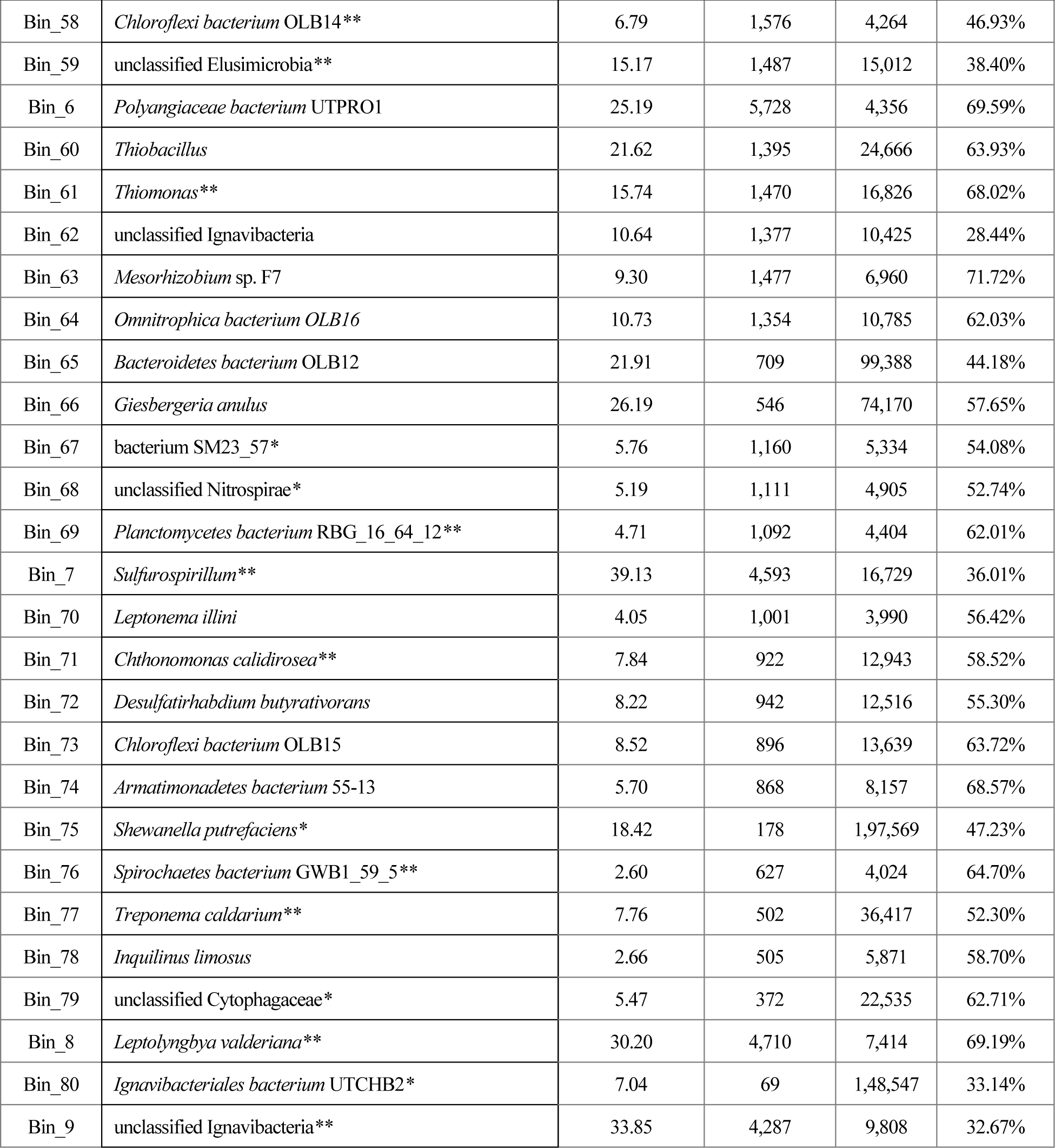
Bins reconstructed from the cross-assembly of three habitats (ecosystem core metagenomics) arev shown here as (*HSG, **ECG)

To investigate habitat specific genes that how these genes are distributed across microbial populations at distinct habitats within the hot spring. Firstly, the geometric and biochemical homogeneity and homogeneity of 39 bacterial MAGs belonged to 12 different phyla were estimated by microbial pangenomic analysis. As most of the MAGs are identically different so, to study the niche specific gene complexity we optimize them together corresponding to their habitat. We identified very complex clade patterning amongst the same phyla candidates and retrieved most explicit gene clusters present in all recovered genomes. In this pangenome, 105,589 overall gene content was partitioned into 15.2% of the maximum occurred gene clusters revealing that (N= 15,982) there are similar bacterial marker (>97% similarity) or essential machinery core-gene repertoire present in MAGs which may be involved in common functional processes required for survival in similar environment (Figure 4). But to define specific niche variation we took 89,607 accessory gene clusters (84.8 % of total pangenome) which were further restricted with rigorous algorithms in order to detect strain specific variability of every MAG which were later called as Habitat Specific Genotypes (HSGs). The HS_g recovered from HSGs were ∼20,676 singleton accessory gene clusters partitioned as n= 11,980 in microbial mat (HS_gM, 1036 ± 25.3), n=773 in sediment (HS_gS, 275 ± 92.8), and n=7,923 in water (HS_gW, 593.4 ± 45), genotypes. Overall, the higher number of HS_g in microbial mat could be explained as diverse group of MAGs were reconstructed from microbial mat than water whereas sediment with only 4 MAGs (not observed as per genome variation) showed lower values.

Secondly, comparative metagenome clustering achieved with ecosystem core metagenomic approach was well appropriate in order to estimate higher number of gene clusters in three habitats. As shown in figure 4B and Table 1, we retrieved a total of ∼55,747 gene clusters in 23 bins partitioned into microbial mat (n= 14,230; 5), sediment (n= 27,731; 12) and water (n= 13,786; 6). We suggested that all the bins derived, have acquired the genetic variations through a series of events with course of time and could be distinct in each taxa (specific) to survive in the microenvironment. The HS_g and HSG_g were further recruited with COG subsystem and biochemical pathways to investigate the distribution of habitat-specific traits across hot spring.

### Biogeochemical cycle inferences using Ecosystem Core Genotypes (ECGs)

Besides identifying the ecosystem core-gene clusters (EC_g), it was necessary to detect their role within very common metabolic potential with emphasis on essential and remarkable processes in this terrestrial ecosystem. A combined metabolic significance of three habitats was determined for very important and key regulatory elements involved in biogeochemical cycles. In this case, the ORF’s of ECGs were blast hit with Pfam domains of biogeochemical pathways of Nitrogen (N), Sulfur (S), Iron (Fe), Oxygen (O) and Carbon (C). The analysis of estimation of biogeochemical cycles gained complete insights in the complex metabolic chemical reactions of nitrogen, sulfur, iron and oxygen cycles in ECGs specified despite their prodigious capacity of molecular transformation. The essential pathways like sulfur reduction (QmoABC), SA reduction (hydACD), ammonia assimilation, superpathway ammonia oxidation, sulfoacetaldehyde degradation (SafD, isfD), thiosulfate disproportionation (Rhodonase) were completed in ECG bins and nitrate reduction, tetrathionate reduction, sulfur assimilation (cys ACDJNPQU), taurine degradation were >60% reconstructed (Figure 5A). The enzyme sulfoactealdehyde belong to short-chain dehydrogenase/reductase (SDR) family was highly abundant which participates in metabolism of taurine and hypotaurine that produces hydrogen sulfide as byproduct (Weinitschke 2010, Rohwerder, 2020). The common metabolic potentital formed in all three habitats revealed a composite activity of utilization of inorganic sulfur (sulfate, thiosulfate, sulfide) and nitrogen (nitrate, ammonia) for energy and survival (LeKieffre *et al.,* 2018). Also, we observed that there is overall abundance of nitrogen cycle with mean entropy (H’) scores of 0.61 followed by Fe (0.58), O (0.41), S (0.31) and C (0.1) (Figure 5B). The scores suggested that there are abundances of nitrogen-fixing microbes and sulfur utilizing bacteria with iron as essential component for physiological processes like replication, transcription, metabolism, and energy generation via respiration.

**Figure 5:**
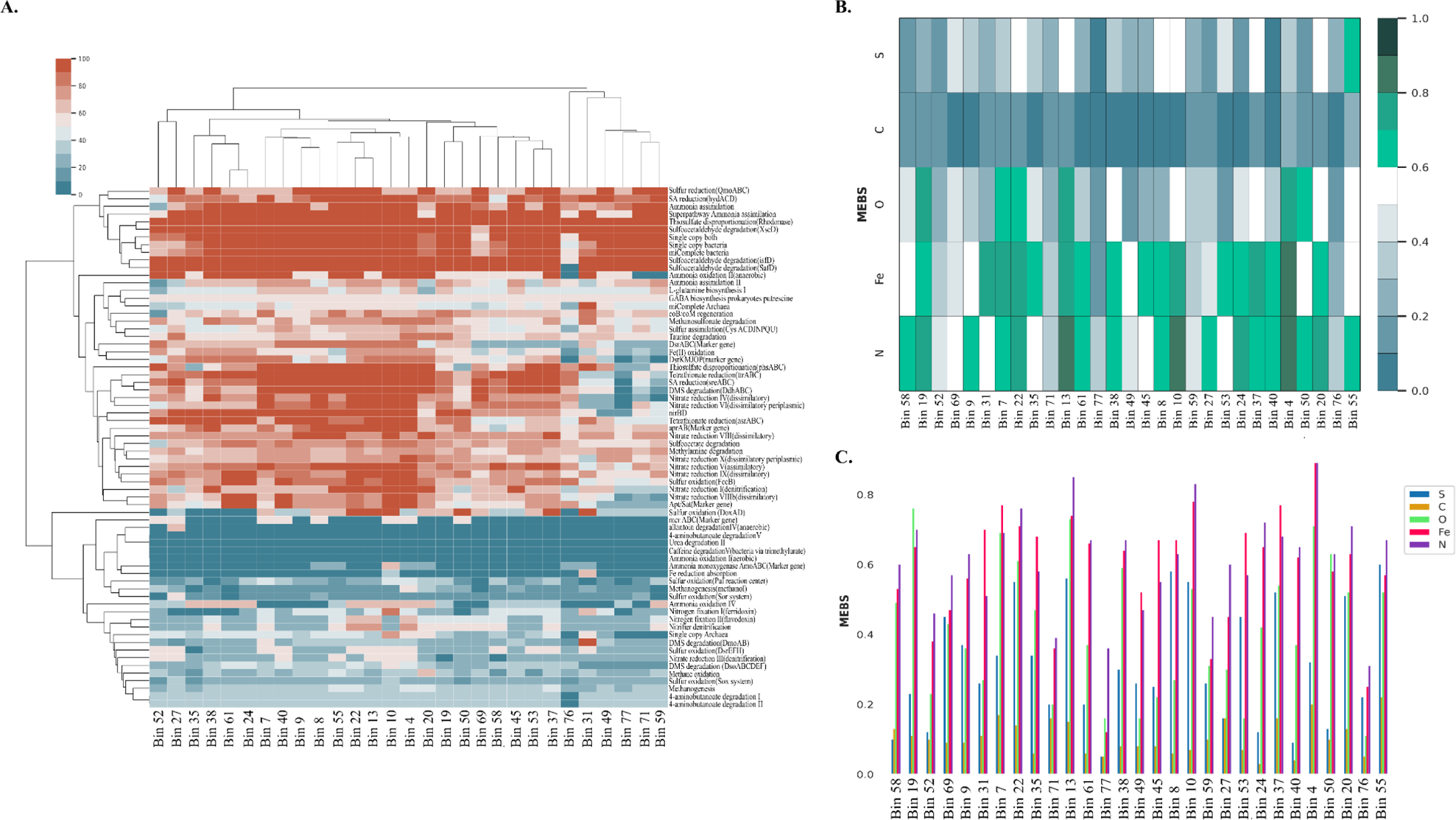
Biogeochemical cycling within the 29 bins reconstructed from orthologous segments of three habitats. (A) Metabolic completeness of essential biogeochemical pathways shown with color gradient, the completed pathways shown in red color and minimal reconstructed in blue color, (B) The entropy scores (H’) of N, S, C, O and Fe were estimated across all MAGs and positioning of MAGs with a lower to higher sulfur entropies in order to represent a relative biogeochemical cycling of microbes in different habitat, (C) Comparative depiction of distribution of biogeochemical cycling in each individual bin.

### Differential functional features of Habitat Specific Genotypes

Based on sequential nucleotide variations detection, achieved through *denovo* strain-level resolution within reconstructed genomes, the habitat specific gene content in individual habitats were annotated to decipher functional attributes. The COGs functional categories reconstructed in HS_g were estimated and sequence profiles of HS_gM, HS_gS and HS_gW showed significant differences. Habitat specific level functional divergence (diverse sequence identical functions) of essential biogeochemical pathways was also evident in all three habitats (Figure 5). On the basis of relative enrichment of protein families, corresponding COG functions and biogeochemical pathways, we were able to resolve the habitat specific quantitative traits with inconstant values (Figure 6). Approximately, 9.5% and 13% of gene clusters corresponded highly to amino acid transport and metabolism (E) among microbial mat and water, respectively while 19.8% of sediment sequences annotated as translation, ribosomal structure and biogenesis (J). The ORFs were also belonged to second abundant energy production and conversion (C) in microbial mat, both J and E in sediment and water, respectively (Figure 6A). Also, the sub-class of proteins that were enriched in habitat specific genetic repertoire majorly classified into ribosomal structures, ABC-type amino acids, translational factors (n >450 in microbial mat), DNA replication and repair mechanism, ATP-dependent proteases. The genetic variations here are directly revealing the adaptation of microbes under three niches. The single nuclueotide polymorphism (SNPs), mutation, heterogeneity, functional diversification, lateral gene transfer (LGT) and speciation are the major events occurred within the species accounted for the habitat specific changes (reference). Here, we suggested that the HS_g are homologous set of functional genes essential for HSGs survival in the identical environmental conitions such as low pH, surface temperature, metal or ion contamination with different thresholds.

**Figure 6:**
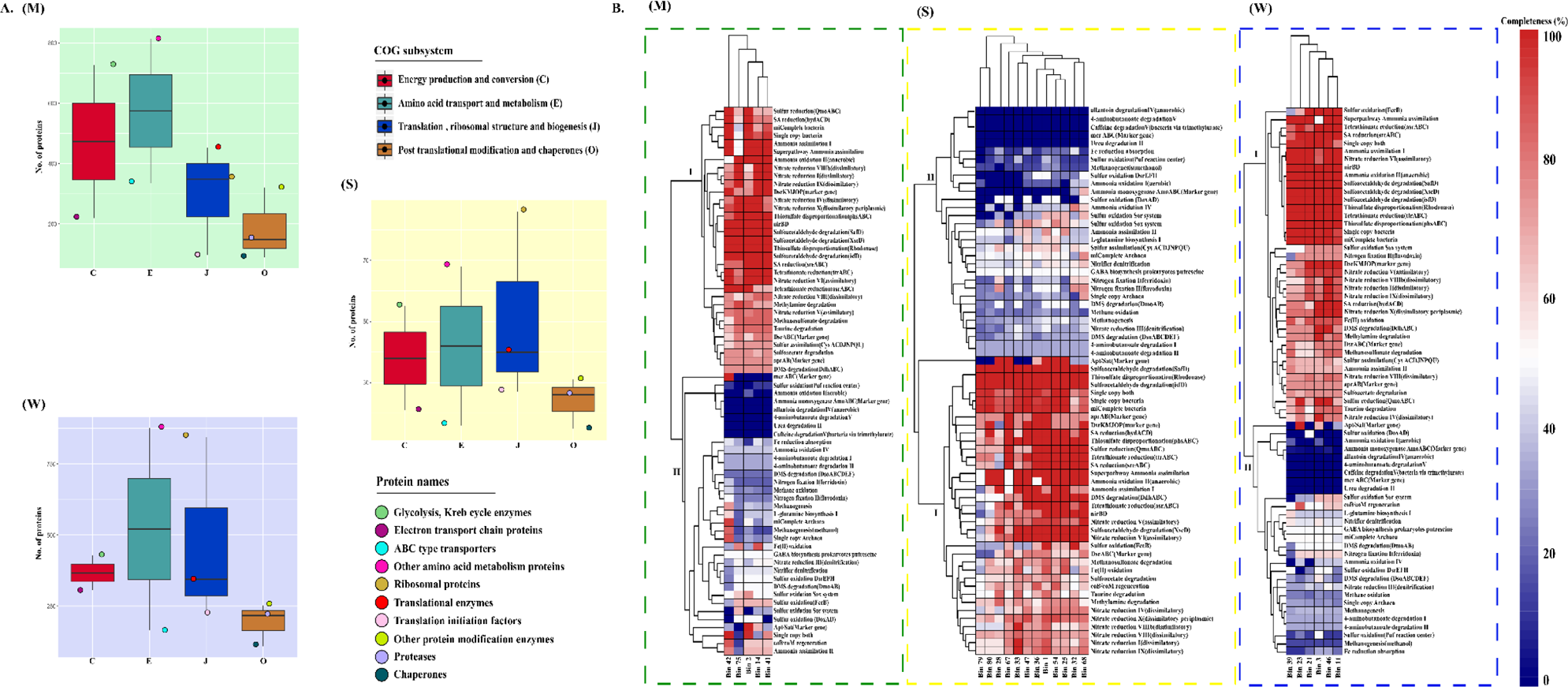
Functional attributes of habitat-specifc gene clusters identified using COG subsystem classification and biogeochemical cycle pathways completeness. A) the classification of HS_g into COG subsystem classes with their abundant proteins and (B) estimation of completeness of metabolic cycling in HSG_g, and represented in all three habitats where (M) as microbial mat, (S) as sediment, (W) as water.

As, the genetic repertoire of these diverse bins vary according to their speciation, functional diversification and lateral gene transfer, we concluded that the chances of survival of these bacteria are equal but the alteration in their genetic makeup is differential. In this case, we observed that the maximum of HS_g were not orthologs but encoding for same metabolic properties which is projected to be varied according to the adaptability of the microorganism (Pacciani-Mori *et al.,* 2020). So, the strain-level resolution categorised in COG subsystem revealed only major and general fundamentals of habitats. Therefore, we tried to reveal the most distinctive features that were acquired by the microbiota in individual habitats by comparing the metagenomes (HSGs). Although, the configuration of microenvironment in habitats using both strain level resolution and ecosystem core metagenomics were ascertain. After generating HSGs (Figure 1c), their metabolism pathways were clustered together at a stringent dissimilarity method using hclust (“complete”) Euclidean distance approach method in which we categorised them as clade I (>60%, enrichment) and clade II (minimum relative abundance) (Figure 6B). In clade I, we observed that the maximum abundant metabolic pathways in all three habitats were nitrate reduction (dissimilatory), superpathway ammonia oxidation, tetrathionate reduction, sulfoacetaldehyde degradation (SafD, isfD), thiosulfate disproportionation (Rhodonase), nirBD, SA reduction. Interestingly, genetic enrichment for coB/coM regeneration pathway was only observed (>63% reconstruction) across HSGs of sediment whereas only HSGs of water were found enriched for sulfur oxidation (FccB), ammonia assimilation II, nitrogen fixation II (flavodoxin), sulfur oxidation (sox) (Nagar et al., 2020).

## Conclusions

Here, we chose the meso-thermic habitats of an equivalent hot spring ranged temperature (42 – 58℃) to relate the core and habitat-specific attributes. In conclusion, our results reveal the functional potential of habitats and detailed association of microbes between or within the habitat using habitat-habitat interactions, genome-resolved metagenomics and ecosystem core metagenomics. Using these omics modification we were able to uncover the predominant microbes and gene families involved in energy conservation and biogeochemical cycling. In the case, we observed that the generalist and specialist species have differential abundance of taxa but executing similar functions with altered genetic repertoire. The approach delivers brief insights in shared and specialised genetic repertoire and explores the functional dynamics at different saptio-temporal scales, targeting microbial communities from separated habitats, along with similar environmental gradients. The HSGs were altering according to the complexity and there was also an inter-specific competition with other members present in the same habitat facilitating them to retain large-scale homologous gene clusters. Ecosystem core metagenomics studies have classified the basis of linkages between ECGs and their interrelation while being distributed in different habitats.

## Acnowledgement

This work was supported by funds from the Department of Biotechnology (DBT), National Bureau of Agriculturally Important Microorganisms (NBAIM), University Grants Commission-Career Advancement Scheme and Department of Science and Technology-Purse grant. S.N. and C.T. thank Council of Scientific and Industrial Research (CSIR) for providing doctoral fellowships.

## Author Contributions

SN and RKN planned the study. SN and CT performed the analysis. SN and CT wrote the manuscript. RKN critically reviewed the manuscript and improved it. All authors read and approved the final manuscript

## References

1. Langmead B, Salzberg S: Fast gapped-read alignment with Bowtie 2. Nat Methods 2012, 9:357–359.

2. Bolan NS, Hedley MJ: Role of carbon, nitrogen, and sulfur cycles in soil acidification. In Handbook of Soil Acidity (CRC Press, 2003).

3. Bowen de León K, Gerlach R, Peyton BM, Fields MW: Archaeal and bacterial communities in three alkaline hot springs in Heart Lake Geyser Basin, Yellowstone National Park. Front Microbiol 2013, 4:330.

4. Caporaso JG, Kuczynski J, Stombaugh J, Bittinger K, Bushman FD, Costello EK, Fierer N, Peña AG, Goodrich JK, Gordon JI, Huttley GA, Kelley ST, Knights D, Koenig JE, Ley RE, Lozupone CA, McDonald D, Muegge BD, Pirrung M, Reeder J, Sevinsky JR, Turnbaugh PJ, Walters WA, Widmann J, Yatsunenko T, Zaneveld J, Knight R: QIIME allows analysis of high-throughput community sequencing data. Nat Methods 2010, 7:335–6.

5. Li D, Liu CM, Luo R, Sadakane K, Lam TW: MEGAHIT: an ultra-fast single-node solution for large and complex metagenomics assembly via succinct de Bruijn graph Bioinformatics 2015, 31:1674–1676

6. de la Torre JR, Walker CB, Ingalls AE, Konneke M, Stahl DA: Cultivation of a thermophilic ammonia oxidizing archaeon synthesizing crenarchaeol. Environ Microbiol 2008, 10:810–818.

7. Eme L, Reigstad LJ, Spang A, Lanzén A, Weinmaier T, Rattei T, Schleper C, Brochier-Armanet C: Metagenomics of Kamchatkan hot spring filaments reveal two new major (hyper)thermophilic lineages related to Thaumarchaeota. Res Microbiol 2013, 164:425–38.

8. Eme L, Reigstad LJ, Spang A, Lanzén A, Weinmaierf T, Rattei T, Schleper C, Brochier-Armanet C: Metagenomics of Kamchatkan hot spring filaments reveal two new major (hyper) thermophilic lineages related to Thaumarchaeota. Res Microbiol 2013, 164:425e438.

9. Kopylova E, Noé L, Touzet H: SortMeRNA: fast and accurate filtering of ribosomal RNAs in metatranscriptomic data. Bioinformatics 2012, 28(4):3211–3217.

10. Foster ZSL, Sharpton TJ, Grünwald NJ: Metacoder: An R package for visualization and manipulation of community taxonomic diversity data. PLoS Comput Biol 2017, 13: e1005404.

11. Galperin MY, Kristensen DM, Makarova KS, Wolf YI, Koonin EV: Microbial genome analysis: the COG approach. Brief Bioinform. 2019, 20(4):1063–1070.

12. Hall JR, Mitchell KR, Jackson-Weaver O, Kooser AS, Cron BR, Crossey LJ et al: Molecular characterization of the diversity and distribution of a thermal spring microbial community by using rRNA and metabolic genes. Appl Environ Microbiol 2008, 74:4910–4922.

13. Hatzenpichler R, Lebedeva EV, Spieck E, Stoecker K, Richter A, Daims H, Wagner M: A moderately thermophilic ammonia-oxidizing crenarchaeote from a hot spring. Proc Natl Acad Sci USA 2008, 105(6):2134–2139.

14. Huang Y, Niu B, Gao Y, Fu L, Li W: CD-HIT Suite: a web server for clustering and comparing biological sequences. Bioinformatics 2010, 26, 680–682.

15. Husson O: Redox potential (Eh) and pH as drivers of soil/plant/microorganism systems: a transdisciplinary overview pointing to integrative opportunities for agronomy. Plant Soil 2013, 362:389–417 (2013).

16. IBM Corp. Released: IBM SPSS Statistics for Windows, Version 25.0. Armonk 2017, NY:IBM Corp.

17. Inskeep WP, Jay ZJ, Tringe SG, Herrgard MJ, Rusch DB: The YNP metagenome project: environmental parameters responsible for microbial distribution in the Yellowstone geothermal ecosystem Front Microbiol 2013, 4:10.3389

18. Inskeep WP, Rusch DB, Jay ZJ, Herrgard MJ, Kozubal MA, Richardson TH et al: Metagenomes from High-Temperature Chemotrophic Systems Reveal Geochemical Controls on Microbial Community Structure and Function. PLoS ONE 2010, 5:e9773.

19. Alneberg J, Bjarnason B, de Bruijn I et al: Binning metagenomic contigs by coverage and composition. Nat Methods 2014, 11:1144–1146.

20. Kanehisa M, Goto S, Kawashima S, Okuno Y, Hattori M (2004) The KEGG resource for deciphering the genome. Nucleic Acids Res 2004, 32: 277–280.

21. Kang D, Li F, Kirton ES, Thomas A, Egan RS, An H, Wang Z: MetaBAT 2: an adaptive binning algorithm for robust and efficient genome reconstruction from metagenome assemblies. PeerJ Preprints 2019, 7:e27522v1

22. Kim D, Song L, Breitwieser FP, and Salzberg SL: Centrifuge: rapid and sensitive classification of metagenomic sequences. Genome Res 2016, (12):1721–1729

23. Koch T, Dahl C: A novel bacterial sulfur oxidation pathway provides a new link between the cycles of organic and inorganic sulfur compounds. ISME J. 2018, 12:2479–2491.

24. Kolde R, Kolde MR: Package ‘pheatmap’. (2015) https://cran.rproject.org/web/packages/pheatmap/pheatmap.pdf

25. Lehmann A, Zheng W, Rillig MC: Soil biota contributions to soil aggregation. Nat Ecol Evol 2017, 1:1828–1835.

26. LeKieffre, C., Jauffrais, T., Geslin, E., et al: Inorganic carbon and nitrogen assimilation in cellular compartments of a benthic kleptoplastic foraminifer. Sci Rep 2018, 8:10140.

27. Letunic I, Bork P: Interactive tree of life (iTOL) v3: an online tool for the display and annotation of phylogenetic and other trees. Nucleic Acids Res 2016, 44:W242–W245.

28. Li H, Handsaker B, Wysoker A, Fennell T, Ruan J, Homer N, Marth G, Abecasis G, Durbin R, 1000 Genome Project Data Processing Subgroup: The Sequence alignment/map (SAM) format and SAMtools, Bioinformatics 2009, 25(16):2078–9

29. Lozupone CA, Knight R: Global patterns in bacterial diversity. Proc Natl Acad Sci USA 2007, 104:11436–11440.

30. Ma ZS, Guan Q, Ye C: Network analysis suggests a potentially ‘evil’ alliance of opportunistic pathogens inhibited by a cooperative network in human milk Bacterial communities. Sci Rep 2015, 5:8275.

31. Ma ZS, Li L, Li W, et al: Integrated network-diversity analyses suggest suppressive effect of Hodgkin’s lymphoma and slightly relieving effect of chemotherapy on human milk microbiome. Sci Rep 2016, 6:28048.

32. Meyer F, Paarmann D, D’Souza M, Olson R, Glass EM, Kubal M, Paczian T, Rodriguez A, Stevens R, Wilke A, Wilkening J, Edwards RA: The metagenomics RAST server - a public resource for the automatic phylogenetic and functional analysis of metagenomes. BMC Bioinformatics, 2008, 9:386.

33. Meyer-Dombard DR, Shock EL, Amend JP: Archaeal and bacterial communities in geochemically diverse hot springs of Yellowstone National Park, USA. Geobiology 2005, 3:211–227.

34. Nagar S, Talwar C, Haider S, Puri A, Ponnusamy K, Gupta M, Sood U, Bajaj A, Lal R and Kumar R: Phylogenetic Relationships and Potential Functional Attributes of the Genus *Parapedobacter*: A Member of Family *Sphingobacteriaceae*. Front. Microbiol. 2020, 11:1725.

35. Nagar S, Talwar C, Motelica-Heino M, Richnow H-H, Shakarad M, Lal R, Negi RK: Microbial Ecology of Sulfur Biogeochemical Cycling at a Mesothermal Hot Spring Atop Northern Himalayas, India. Front Microbiol 2022, 13:848010.

36. Nagar S, Bharti M, Negi RK: Genome-resolved metagenomics revealed metal-resistance, geochemical cycles in a Himalayan hot spring. Appl Microbiol Biotechnol 2023, 107:3273–3289.

37. Pagaling E, Grant WD, Cowan DA, Jones BE, Ma Y, Ventosa A et al: Bacterial and archaeal diversity in two hot spring microbial mats from the geothermal region of Tengchong, China. Extremophiles 2012, 16:607–618.

38. Parks DH, Imelfort M, Skennerton CT, Hugenholtz P, Tyson GW: Assessing the quality of microbial genomes recovered from isolates, single cells, and metagenomes. Genome Res 2014, 25:1043–1055.

39. Pester M, Schleper C, Wagner M: The Thaumarchaeota: an emerging view of their phylogeny and ecophysiology. Curr Opin Microbiol 2011, 14:300–306.

40. Philippot L, Chenu C, Kappler A et al: The interplay between microbial communities and soil properties. Nat Rev Microbiol 2024, 22:226–239.

41. Eddy SR: Profile Hidden Markov Models. Bioinformatics 1998, 14:755–763.

42. Quast C, Pruesse E, Yilmaz P, Gerken J, Schweer T, Yarza P, Glöckner, FO: The SILVA ribosomal RNA gene database project: improved data processing and web-based tools. Nucleic Acids Res 2013, 41(Database issue):D590–D596.

43. Quince C, Delmont TO, Raguideau S, Alneberg J, Darling AE, Collins G, Eren AM: DESMAN: a new tool for de novo extraction of strains from metagenomes. Genome Biol 2017, 18(1):181.

44. Shannon P, Markiel A, Ozier O, Baliga NS, Wang JT, Ramage D, Amin N, Schwikowski B, Ideker T. (2003) Cytoscape: a software environment for integrated models of biomolecular interaction networks. Genome Res 13:2498–504.

45. Sokol NW et al: Life and death in the soil microbiome: how ecological processes influence biogeochemistry. Nat Rev Microbiol 2022, 20:415–430.

46. Tatusov RL, Fedorova ND, Jackson JD, Jacobs AR, Kiryutin B, Koonin EV: The COG database: an updated version includes eukaryotes. BMC Bioinformatics 2003, 4: 41.

47. Truong DT, Franzosa EA, Tickle TL, Scholz M, Weingart G, Pasolli E, Tett A, Huttenhower C, Segata N: MetaPhlAn2 for enhanced metagenomic taxonomic profiling. Nat Methods 2015. 12(10):902–3.

48. Varin T, Lovejoy C, Jungblut AD, Vincent WF, Corbeil J: Metagenomic profiling of Arctic microbial mat communities as nutrient scavenging and recycling systems. Limnol Oceanogr 2010, 55:1901–1911

49. Wang S, Hou W, Dong H, Jiang H, Huang L, Wu G et al: Control of temperature on microbial community structure in hot springs of the Tibetan Plateau. PLoS ONE 2013, 8:e62901.

50. Whitman WB, Coleman DC, Wiebe WJ: Prokaryotes: the unseen majority. Proc. Natl Acad. Sci. USA 1998, 95:6578–6583.

51. Wickham H: ggplot2: elegant graphics for data analysis. Springer, New York, NY 2009.

52. Odum EP, Smalley AE: Comparison of population energy flow of a herbivorous and a deposit-feeding invertebrate in a salt marsh ecosystem. Proc Natl Acad Sci USA 2009, 45(4):617–2.

53. Dussault AC, Mermans E, Barker G et al:Functional Diversity: An Epistemic Roadmap, BioScience 2019, 69(10):800–811.

54. Wang XB, Lü XT, Yao J et al: Habitat-specific patterns and drivers of bacterial β-diversity in China’s drylands. ISME J 2017, 11:1345–1358.

55. Seto M, Iwasa Y: The fitness of chemotrophs increases when their catabolic by-products are consumed by other species. Ecol Lett 2019, 22(12):1994–2005.

56. Schimel JP, Schaeffer SM: Microbial control over carbon cycling in soil. Front. Microbiol 2012, 3:348.

57. Chu H, Fierer N, Lauber CL, Caporaso JG, Knight R: Soil bacterial diversity in the Arctic is not fundamentally different from that found in other biomes. Environ Microbiol 2010, 12:2998–3006.

58. Maestre FT, Delgado-Baquerizo M, Jeffries TC, Eldridge DJ, Ochoa V, Gozalo B, et al: Increasing aridity reduces soil microbial diversity and abundance in global drylands. Proc Natl Acad Sci 2015, 112:15684–15689.

59. Prober SM, Leff JW, Bates ST, Borer ET, Firn J, et al. 2015. Plant diversity predicts beta but not alpha diversity of soil microbes across grasslands worldwide. Ecol Lett 2015, 18:85–95.

60. Hubbell SP: Neutral theory in community ecology and the hypothesis of functional equivalence. Funct Ecol 2005, 19:166–172.

61. Ronca S, Ramond JB, Jones BE, Seely M, Cowan DA: Namib Desert dune/interdune transects exhibit habitat-specific edaphic bacterial communities. Front Microbiol 2015, 6:12.

62. Yergeau E, Newsham KK, Pearce DA, Kowalchuk GA: Patterns of bacterial diversity across a range of Antarctic terrestrial habitats. Environ Microbiol 2007, 9: 2670–2682

63. Benjamini Y, Hochberg Y: Controlling the false discovery rate: a practical and powerful approach to multiple hypothesis testing. J R Stat Soc B 1995, 57:289–300

64. Zinger L, Boetius A, Ramette A: Bacterial taxa-area and distance-decay relationships in marine environments. Mol Ecol 2014, 23:954–964.

65. Callahan BJ, McMurdie PJ, Rosen MJ, Han AW, Johnson AJ, Holmes SP: DADA2: High-resolution sample inference from Illumina amplicon data. Nat Methods 2016, 13(7):581–3

66. DeSantis TZ, Hugenholtz P, Larsen N, Rojas M, Brodie EL, Keller K, Huber T, Dalevi D, Hu P, Andersen GL: Greengenes, a chimera-checked 16S rRNA gene database and workbench compatible with ARB. Appl Environ Microbiol 2006, 72(7):5069–72.

67. Yuan C, Lei J, Cole J, Sun Y: Reconstructing 16S rRNA genes in metagenomic data. Bioinformatics 2015, 31:i35–i43.

68. Cole JR, Chai B, Farris RJ, Wang Q, Kulam-Syed-Mohideen AS et al: The ribosomal database project (RDP-II): introducing myRDP space and quality controlled public data. Nucleic Acids Res 2007, 35:D169–D172

69. Bolyen E, Rideout JR, Dillon MR, Bokulich NA, Abnet CC: Reproducible, interactive, scalable and extensible microbiome data science using QIIME 2. Nat Biotechnol 2019, 37:852–857.

70. Kumar S, Stecher G, Li M, Knyaz C, Tamura K: MEGA X: molecular evolutionary genetics analysis across computing platforms. Mol Biol Evol 2018, 35:1547

71. McDonald JH: Handbook of biological statistics. Baltimore, MD: sparky house publishing 2009, 2:6–59.

72. Eren AM, Esen ÖC, Quince C, Vineis JH, Morrison HG et al: Anvi’o: an advanced analysis and visualization platform for ‘omics data. PeerJ 2015, 3:e1319.

73. Hyatt D, Chen GL, Locascio PF, Land ML, Larimer FW, Hauser LJ: Prodigal: prokaryotic gene recognition and translation initiation site identification. BMC Bioinform 2010, 11:119

74. Finn RD, Bateman A, Clements J, Coggill P, Eberhardt RY, et al: Pfam: the protein families database. Nucleic Acids Res 2014, 42: D222–30.

75. De Anda V, Zapata-Peñasco I, Poot-Hernandez AC, Eguiarte LE, Contreras-Moreira B, et al: MEBS, a software platform to evaluate large (meta)genomic collections according to their metabolic machinery: unraveling the sulfur cycle. Gigascience 2017, 6:1–17.

76. Rohwerder T: New structural insights into bacterial sulfoacetaldehyde and taurine metabolism. Biochem J 2020, 477:1367–1371.

77. Weinitschke S: New intermediates, pathways, enzymes and genes in the microbial metabolism of organosulfonates (Doctoral dissertation) 2010.

78. Pacciani-Mori L, Giometto A, Suweis S, Maritan A: Dynamic metabolic adaptation can promote species coexistence in competitive microbial communities. PLoS Comput Biol 2020, 16:e1007896.

79. Zhang L, Delgado-Baquerizo M, Shi Y, Liu X, Yang Y, Chu H: Co-existing water and sediment bacteria are driven by contrasting environmental factors across glacier-fed aquatic systems. Water Res 2021, 198:117139.

80. Fray D, McGovern CA, Casamatta DA, Biddanda BA, Hamsher SE: Metabarcoding reveals unique microbial mat communities and evidence of biogeographic influence in low-oxygen, high-sulfur sinkholes and springs. Ecol Evol 2024, 14(3):e11162.

