## Supplementary material for "Integrated multi-platform approaches to gain insights into ecosystem’s fundamental ecology and habitat specific alterations": Supplemntary data

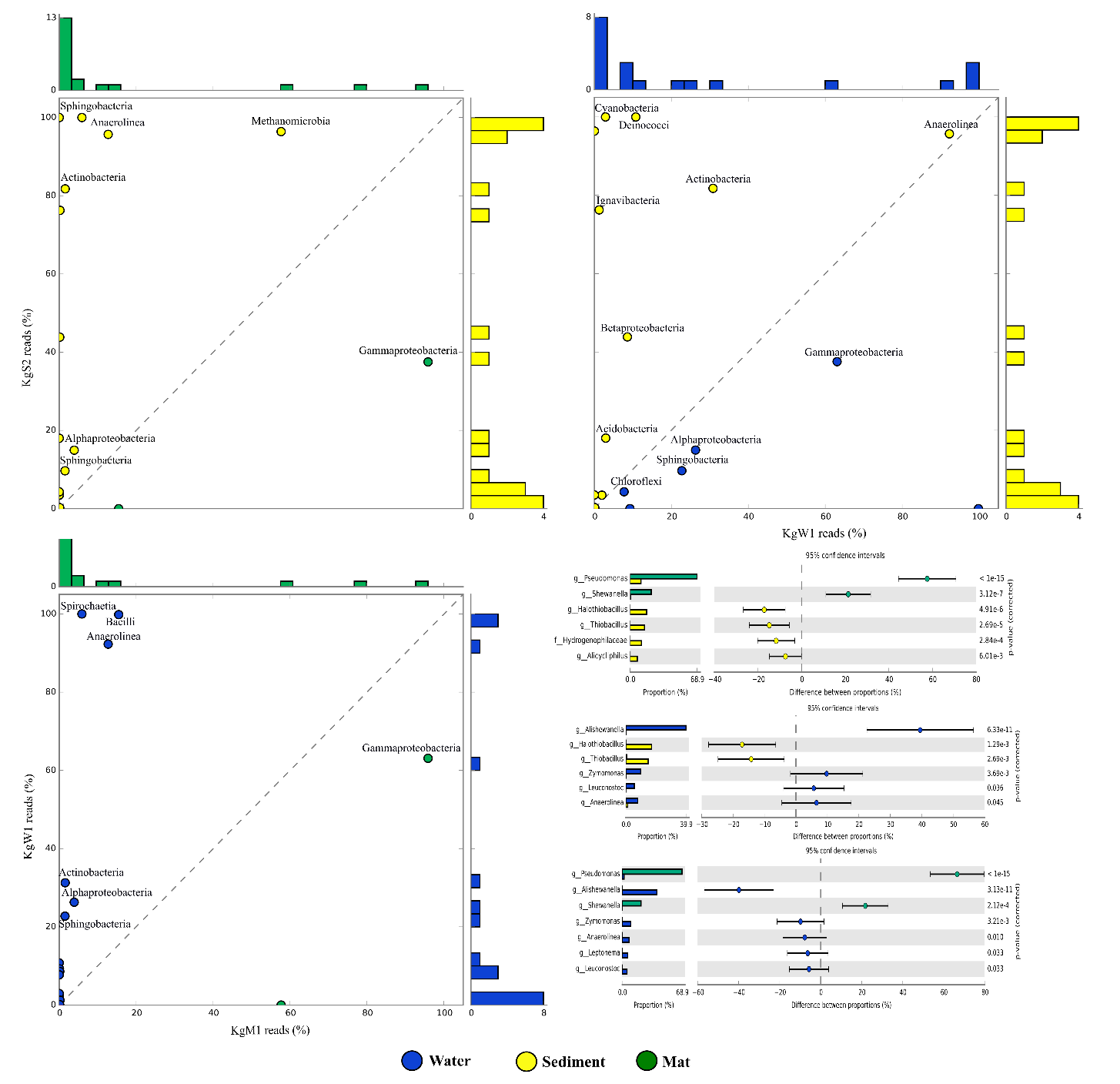

**Figure S1: Correlation of class-level taxonomic profiles between sample types.** (A) Microbial mat and Sediment (R^2^ = 0.542), Water and Sediment (R^2^ = 0.912) Microbial mat and Water (R^2^= 0.711) and genus level taxonomic error plot of dataset vs. dataset at 95% confidence interval.

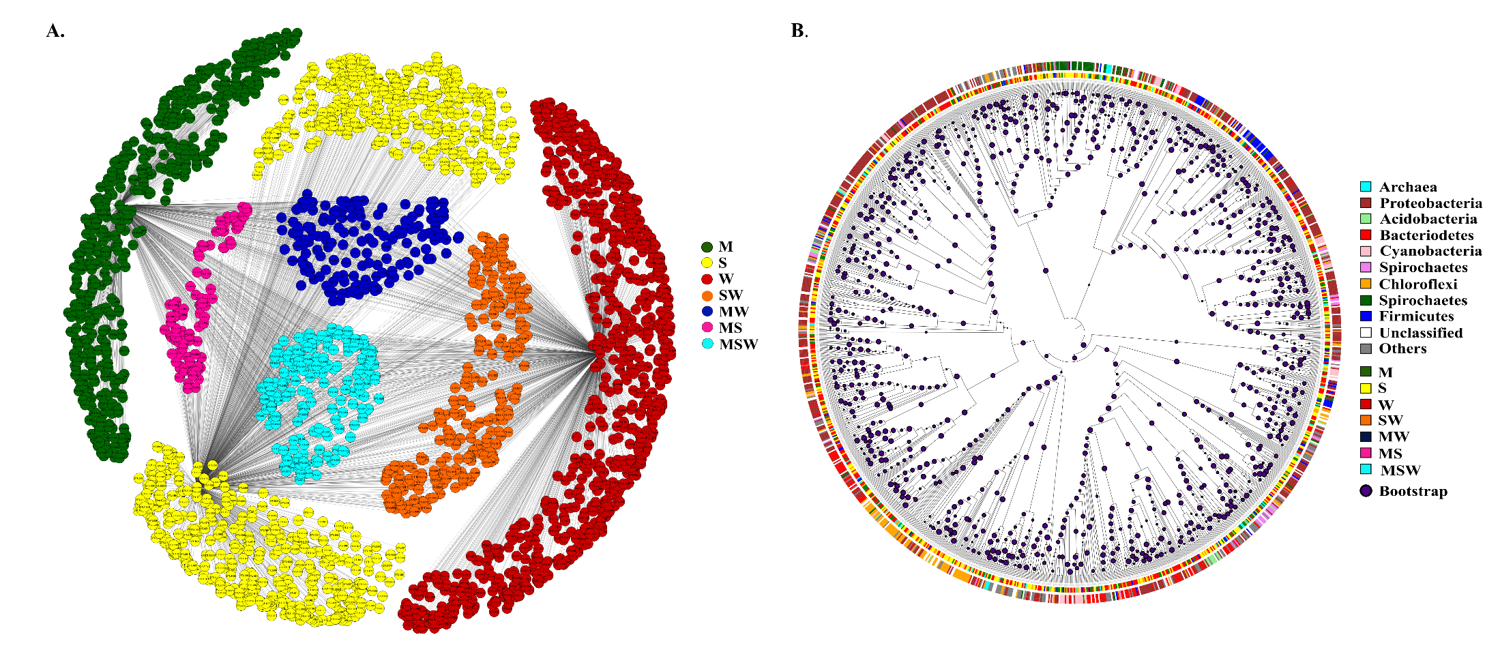

**Figure S2:** The OTUs are aligned together using QIIME2 to form a discrete tree representation. The taxa are assigned according to the 97% similarity to Greengenes database. Inner circle depicting the unique habitats with color green as microbial mat (M), yellow as sediment(S), red as water (W) and common with cyan color (MSW). Purple dots are showing bootstrap greater than 50.
